# Inter-specific Variability in Demographic Processes Affects Abundance-Occupancy Relationships

**DOI:** 10.1101/2020.09.03.282004

**Authors:** Bilgecan Şen, H. Reşit Akçakaya

**Affiliations:** Department of Ecology and Evolution, Stony Brook University, Stony Brook, NY, 11794, U.S.A.

**Keywords:** Abundance, occupancy, density dependence, population demography, intrinsic growth rate, mark-recapture, macroecology

## Abstract

Species with large local abundances tend to occupy more sites. One of the mechanisms proposed to explain this widely reported inter-specific relationship is a cross-scale hypothesis based on dynamics at the population level. Called the vital rates mechanism, it uses within-population demographic processes of population growth and density dependence to predict when inter-specific abundance-occupancy relationships can arise and when these relationships can weaken and even turn negative. Even though the vital rates mechanism is mathematically simple, its predictions has never been tested directly because of the difficulty estimating the demographic parameters involved. Here, using a recently introduced mark-recapture analysis method, we show that there is a weakly positive relationship between abundance and occupancy among 17 bird species. Our results are consistent with the predictions of the vital rate mechanism regarding the demographic processes that are expected to weaken this relationship. Specifically, we find that intrinsic growth rate and local abundance are weakly correlated; and density dependence strength shows considerable variation across species. Variability in density dependence strength is related to variability in species-level local average abundance and intrinsic growth rate; species with lower growth rate have higher abundance and are strongly regulated by density dependent processes, especially acting on survival rates. Species with higher growth rate, on average, have lower abundance and are more weakly regulated by density dependent processes especially acting on fecundity. More generally, our findings support a cross scale mechanism of macroecological abundance-occupancy relationship emerging from density-dependent dynamics at the population level.

## Introduction

Exploring ecological processes and interactions across scales of biological organization allows a deeper understanding of patterns of biodiversity. One well-documented macro-ecological pattern is the abundance-occupancy relationship: across phylogenetically similar species, local average abundance and proportion of sites occupied are positively correlated (Hanski, 1982; Bock & Ricklefs, 1983; Brown, 1984, 1995; Gaston & Blackburn, 2000). This pattern has been observed in a variety of taxa, such as mammals (Blackburn *et al*., 1997), birds (Lacy & Bock, 1986; Blackburn *et al*., 1997), butterflies (Conrad *et al*., 2001), mollusks (Russell & Lindberg, 1988), and plants (Guo *et al*., 2000). A meta-analysis on abundance-occupancy relationships found an overall positive correlation but also reported high variability in the strength of this relationship across species realms (Blackburn *et al*., 2006). Some species groups even show zero or negative correlation between abundance and occupancy (Novosolov *et al*., 2017), which may be the result of an ecological process (Reif *et al*., 2006; Symonds & Johnson, 2006; Ferenc *et al*., 2016; Freeman, 2019), or an artifact of the particular metrics used to represent abundance and occupancy (Wilson, 2008).

Several mechanisms have been proposed to explain the positive relationship between abundance and occupancy (reviewed by Gaston *et al*., 1997 and Borregaard & Rahbek, 2010). Among these, one mechanism, known as “vital rates” is notable for being the only one that explains this macro-ecological relationship with population demography. Proposing the vital rates mechanism, Holt *et al*. (1997) argued that the positive correlation between number of occupied sites and local average abundance across species is a consequence of the relationship between population abundance and intrinsic population growth rate (*r*). If source-sink dynamics are not the primary mode of occurrence, a species exist only in sites with *r* > 0, and higher *r* leads to higher abundances. When *r* is determined by a uniform environmental gradient, any environmental factor that increases *r* will also increase abundance at each site and simultaneously increase the number of occupied sites across the same environmental gradient. Differences in intrinsic growth rates among species will lead to the observed positive inter-specific relationship between abundance and occupancy. While Holt *et al*. (1997) used logistic population growth to explain how intrinsic growth rate and density dependence strength can determine the direction (positive, negative, or no relationship) and strength of the abundance-occupancy relationship, the general assumptions and predictions of vital rates mechanism is not dependent on the specific type of population regulation.

We summarize the predictions of vital rates mechanism under three headings:

### 1 – Relationship between intrinsic growth rate and occupancy among species?

Intrinsic growth rate plays a central role in vital rates mechanism because it is the link between abundance and occupancy. Holt *et al*. (1997). postulates that “Any factor which tends to increase *r* across all sites will simultaneously enlarge the number of sites of occupied, and increase abundance of each occupied site”. So, if there is a positive abundance-occupancy relationship among a group of species and if the vital rates mechanism is the major process that determines this relationship, there should also be a positive association between intrinsic growth rate and occupancy. Alternatively, if intrinsic growth rate and occupancy are not closely related then we would expect a weak abundance-occupancy relationship.

### 2 – Relationship between intrinsic growth rate and abundance among and within species

As stated above, in vital rates mechanism, intrinsic growth rate relates occupancy to abundance. So, if there is positive abundance-occupancy relationship in a group of species, and if the vital rates mechanism is the major process that determines this relationship, populations with higher growth rates should also have higher abundances, and species with higher average growth rates should have higher average abundances. Holt *et al. (1997)* demonstrates this possibility with an alternative parameterization of logistic population growth which does not include carrying capacity as an independent parameter. Instead, equilibrium abundance is determined by intrinsic growth rate and density dependence strength. However, rather than forcing the correlation between intrinsic growth rate and abundance with the structure of a population model, they can be modeled to vary independently. If abundance and intrinsic growth rate do not show positive correlation, we would expect a weak abundance-occupancy relationship according to vital rates mechanism.

### 3 – Variability of density dependence strength among species

Holt *et al*. (1997) hypothesized that “[a] species with small *r* but weak density dependence can obtain enormous local abundances compared to another species with large *r* but intense density dependence”. In this scenario, differences in density dependence strengths breaks the expected positive correlation between abundance and intrinsic growth and weakens the abundance occupancy relationship. Accordingly, if there is a positive abundance-occupancy relationship in a group of species, and if the vital rates mechanism is the major process that determines this relationship, density dependence strength across these species should be similar and should not be correlated with intrinsic growth rate and abundance. However, if density dependence strength is positively correlated with intrinsic growth rate and negatively correlated with abundance (as in the scenario above), we should expect a weaker abundance-occupancy relationship.

Here, we use mark-recapture data to estimate the vital rates and population abundances of 17 bird species, and explicitly test the aforementioned predictions of Holt *et al*. (1997)’s vital rates mechanism. While these predictions have been tested using proxy parameters, for example, the skewness in population size distributions (Freckleton *et al*., 2006), to our knowledge, they have never been explicitly tested with the estimation of the demographic parameters involved in the mechanism itself.

## Materials and Methods

### Data: Mapping Avian Productivity and Survivorship (MAPS)

MAPS is a collaborative mark-recapture program initiated and organized by the Institute for Bird Populations (IBP) since 1989 across 1200 bird banding stations with mist nets in US and Canada. The program provides data for more than 300 species. We used only captures from conterminous United States and from 1992 to 2008 between May and August (breeding season). We considered first year individuals (MAPS age code 2 and 4) as juveniles, and all older individuals (MAPS age code 1,5,6,7, and 8) as adults. Following Albert *et al*. (2016) we only used species with:

1. More than 74 adult individuals, and more than 74 juvenile individuals captured,
2. More than 14 adults, and more than 14 juveniles re-captured.

These filters reduced the dataset to 62 species. We applied CJS-pop to only 42 species which had reasonable data size (e.g. maximum model convergence time was one week). For each species, we treated separate MAPS locations (a cluster of mist-netting and banding stations) to be independent breeding populations. We only included data from populations which have been monitored for at least 5 years.

### Data: Environmental Covariates

We used the downscaled Maurer gridded data (Maurer et al., 2012), which was shown to be superior to several other alternatives in terms of its closeness to observed measurements (Behnke *et al*., 2016), to calculate the ten metrics used as environmental covariates in CJS-pop models: annual mean temperature, mean diurnal range, mean temperature of wettest quarter, annual precipitation, precipitation of Warmest Quarter, maximum consecutive dry days, maximum consecutive 5-day precipitation total for March, standard deviation in daily temperatures for March, total precipitation in May, and number of dry days in April. We used annual aggregates of these metrics when building CJS-pop models. Each MAPS location was assigned the covariate values of the grid cell it was located in. If the mist-netting stations of a location was distributed across multiple grid cells we assigned the average value of the covariates across those grid cells to that location. We repeated this process for every year between 1992-2008 for every MAPS location, creating a time series of weather variables at each location.

### Mark-recapture Analysis Framework

We used the framework developed by Ryu *et al*. (2016), which employs robust design mark-recapture data, to estimate population vital rates (survival and fecundity) as well as density-dependence and process variance in these vital rates (temporal or spatial variability). Models were fit using Bayesian statistics and JAGS as the MCMC sampler. Below, we focus on how these vital rates and capture probabilities are used to estimate intrinsic growth rate and abundance. A more detailed explanation of the framework can be found in Appendix S2.

We model survival and fecundity as functions of environmental covariates in addition to density dependence. An example model structure, here illustrated for *ϕ*_*x,k,t*_ the survival probability of a stage *x* invidiual, at population *k*, and at year *t*, with two environmental covariates is given as:

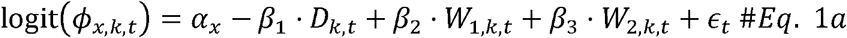

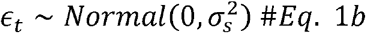

where *α*_*x*_ is the survival probability of a stage *x* individual in logit scale at mean population size and at mean of environmental covariates *W*_1_ and *W*_2_: *β*_1_ is the change in survival in logit scale with one unit change in population density index; *D* is the population density index at population *k*, and at year t (See appendix S2 for density index calculation); *β*_*2*_ and *β*_3_ are the change in survival in logit scale with one unit change in *W*_1_ and *W*_2_, respectively; *ϵ*_*t*_ is the temporal random effect at year *t*, and 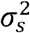 is the temporal process variance of survival at logit scale; *x* = 1,2, …, *X; k* = 1,2,3…, *K; t* =1,2,3…, *T*. We denote *x =* 1 as juveniles and, *x =* 2 as adults.

Similarly, fecundity *F*_*kt*_ at population *k* and year *t* can be modelled as:

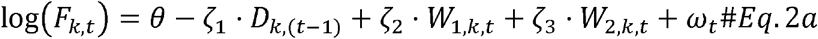

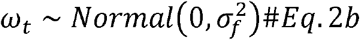

where *θ* is the average fecundity in log scale at mean population size and at mean of environmental covariates *W*_1_ and *W*_2_; *ζ*_1_ is the change in fecundity in log scale with one unit change in population density index; *ζ*_2_ and *ζ*_3_ are the changes in fecundity in log scale with one unit change in *W*_1_ and *W*_2_, respectively; *ω*_*t*_ is the temporal random effect at year *t*, and 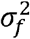 is the temporal process variance of fecundity in log scale. This framework is set so that when a population is at its average density index value (e.g. at carrying capacity), the growth rate calculated by fecundity, adult survival, and juvenile survival is 0 (See appendix S2 for details).

Because this framework is an extended version of a typical Cormack-Jolly-Seber (CJS) method (Ryu *et al*., 2016), capture probability *p*_*x,k,t,h*_ of a stage x individual at population k, year t, and month h is explicitly modelled alongside vital rates as a function of capture effort:

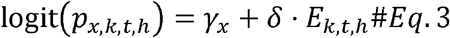

where *γ*_*x*_ is the monthly capture probability on logit scale of a stage x individual at mean capture effort; *δ* is the change in monthly capture probability on logit scale with one unit change in capture effort; E is the capture effort, which is calculated as total mist netting hours per month in a population; *h* = 1,2,3…,*H*.

To prevent overfitting and fasten the convergence of CJS-pop, we reduced the number of covariates with principal component analysis (PCA) and used the number of principle components that cumulatively explains 80% of the variation. We applied PCA separately to each species using the time series of weather covariates from the MAPS locations a given species was captured in.

### Estimating Demographic Parameters

#### Intrinsic Growth Rate

Intrinsic population growth rate is the density independent population growth rate. While the exact meaning depends on the estimation method (Lynch & Fagan, 2009), it represents the theoretical maximum of a population’s ability to grow in size when population density is close to 0. To estimate intrinsic growth rate at a given population and year, we set density to 0 and estimate survival and fecundity only as functions of environmental covariates. We use posterior means of parameters and omit process variance when estimating survival and fecundity. Then, separately for each species, intrinsic growth rate at population *k* and year t can be estimated as:

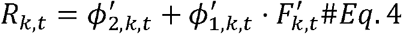

where 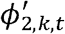 is the adult survival at population *k* and year *t* at 0 density; 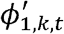 is juvenile survival at 0 density; and 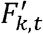 is fecundity at 0 density. We estimate average intrinsic growth rate of each population as geometric mean on log scale:

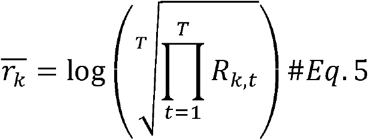

Finally, to estimate species level intrinsic growth rate we use the median of population level growth rates:

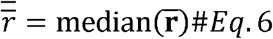

where, 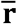 is a vector with population level intrinsic growth rates.

We only used species with a negative density-dependence effect on growth rate (growth rate at density index 2 was lower than growth rate at density index 0). Intrinsic growth rate is not defined for circumstances in which increasing density also increases the population growth; in such a case intrinsic growth rate loses its meaning as a theoretical maximum. Additionally, positive density dependence effects are biologically meaningful only when Allee effects are considered. Because we are not modelling Allee effects with this framework, we removed species that showed a positive density dependence effect on growth rate from further analysis.

#### Population Size

The monthly capture probabilities obtained from CJS-pop are used to calculate the yearly capture probabilities, and these in turn are used to estimate the expected size of each population in each year. Yearly capture probabilities for each adult and juvenile in a given year and population is calculated as follows:

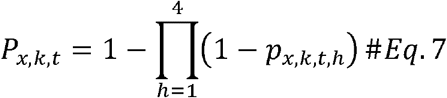

where, *P*_*x,k,t*_ is the probability that a stage *x* individual will be captured at least once in 4 months of the breeding period at year *t* and population *k*.

Using the heuristic estimator for population size with a correction for years with 0 captures (Dail & Madsen, 2011), the numbers of adults and juveniles can be derived as:

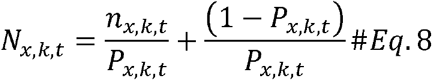

where, *n*_*x,k,t*_ is the number of captured stage *x* individuals at population *k* and year *t*, and *N*_*x,k,t*_ is the expected number of stage *x* individuals of the same population and year combination. Average total population size for each population is estimated as:

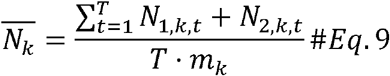

where, *T* is the number of sampling years of a population. MAPS locations can have different number of stations, we standardize total population size by dividing by the number of stations of population *k, m*_*k*_. We estimate the species level population abundance as the median population size across all populations:

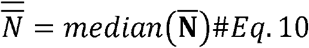

where, 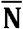 is a vector with average population sizes.

#### Occupancy

There are 495 MAPS locations across US in the dataset we used for this analysis. The number of MAPS locations a species was captured in provides information about the distribution and occupancy of a species. Different measures of occupancy affect the strength of estimated abundance-occupancy relationship (Borregaard & Rahbek, 2010). Here, we define occupancy using two different measures: 1) geographic extent calculated as 100% minimum convex polygon of all the stations in which a species was captured; 2) number of different populations (MAPS locations) in which a species was captured.

### Testing the Predictions of Vital Rates Mechanism

For each test below we estimate the Pearson’s correlation coefficient (*ρ*) and report the posterior probability of the prediction being tested, *P(ρ* > 0):

#### Abundance-occupancy relationship

Between species-level median abundance 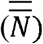 and each of the two metrics of occupancy, across species.

#### Relationship between intrinsic growth rate and occupancy

Between species-level median intrinsic growth rate 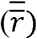 and each of the two metrics of occupancy, across species.

#### Relationship between intrinsic growth rate and abundance within species

Between population-level abundance 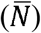 and population level intrinsic growth rate 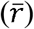, across populations, separately for each species.

#### Relationship between intrinsic growth rate and abundance among species

Between species-level median intrinsic growth rate 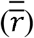 and species-level median abundance 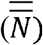, across species.

#### Variability of density dependence strength among species

Among density dependence strength in both survival and fecundity (β and *ζ*, respectively), species-level average intrinsic growth rate 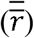, and species-level average abundance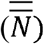.

Because we use uniform priors on *ρ* (Uniformf −1,1)), posterior probability estimates above or below 0.5 can be considered as evidence obtained from data regarding the prediction being tested. Detailed information on priors and model specifications of all Bayesian models used in this analysis can be found in Appendix S2. R code for data manipulation and JAGS code for the mark-recapture framework are available at https://github.com/bilgecansen/VitalRatesTest. Results of mark-recapture models and data for testing the predictions of vital rates mechanism is in Data S1.

## Results

Of the 42 species analyzed, 34 yielded posterior distributions that converged (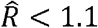 for all parameters). Of these 34 species, only 17 had an overall negative density dependence effect on growth rate (Table 1). Among these 17 species, population size showed a right skewed distribution with many small and few large populations (Appendix S1: Fig. S1a). The majority of these populations have 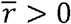 with unimodal distribution that is centered on 0.14 (Appendix S1: Fig. S1b).

**Table 1.**
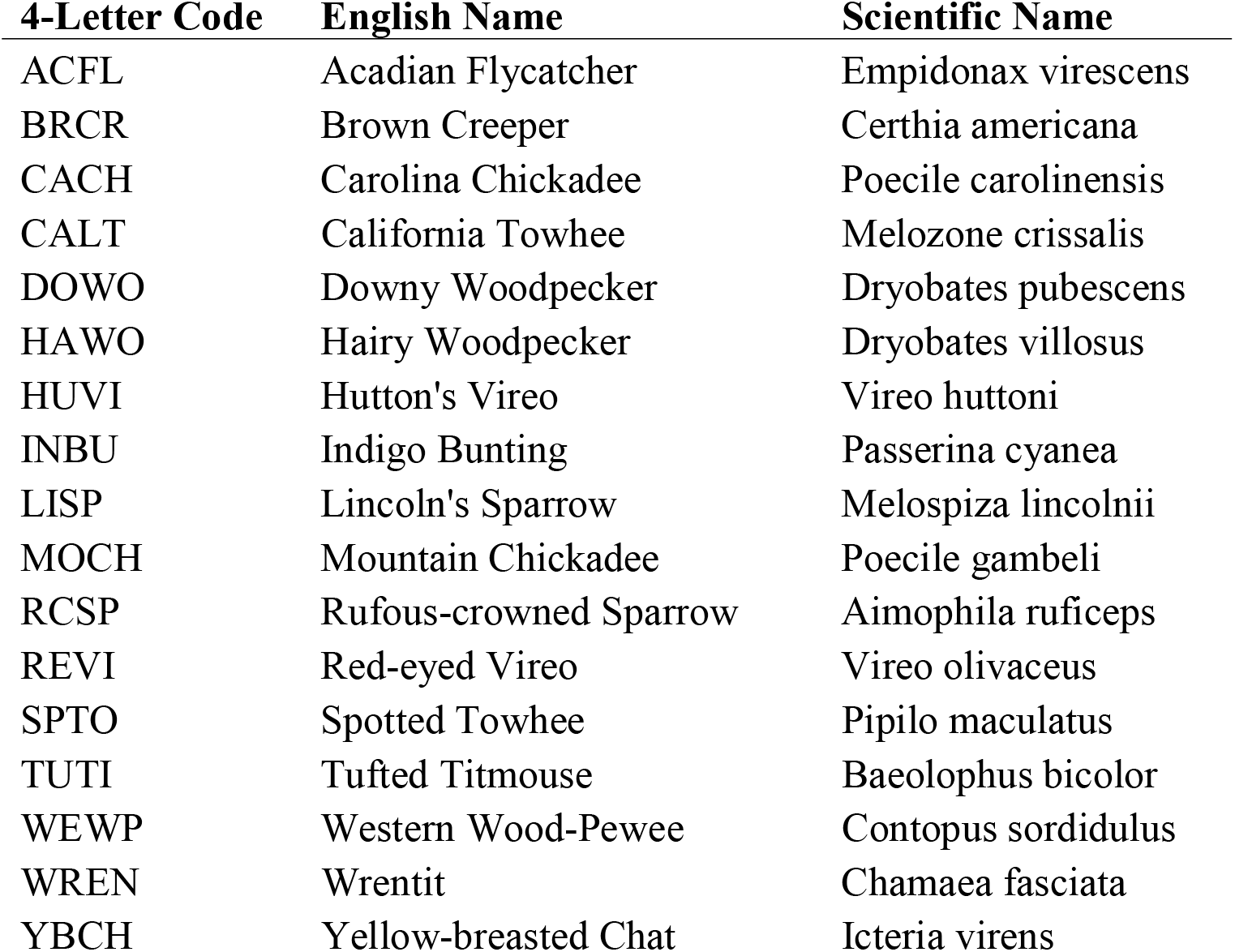
Names of the17 selected species to be used in testing the vital rates mechanism.

We found no evidence that species level abundance 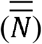 is correlated with geographic extent *(ρ* = 0.012, *P(ρ* > 0) = 0.52; Fig. 1a). There is, however, a weakly positive relationship between 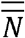 and number of populations *(ρ =* 0.26, *P(ρ* > 0) = 0.86; Fig. 1b). We detected a weakly positive relationship between species level intrinsic growth rate 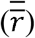 and geographic extent *(ρ =* 0.13, *P(ρ* > 0) = 0.70; Fig. 2a), and a weakly negative relationship between 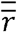 and number of populations *(ρ =* −0.10, *P*(*ρ* > 0) = 0.35; Fig. 2b).

**Figure 1.**
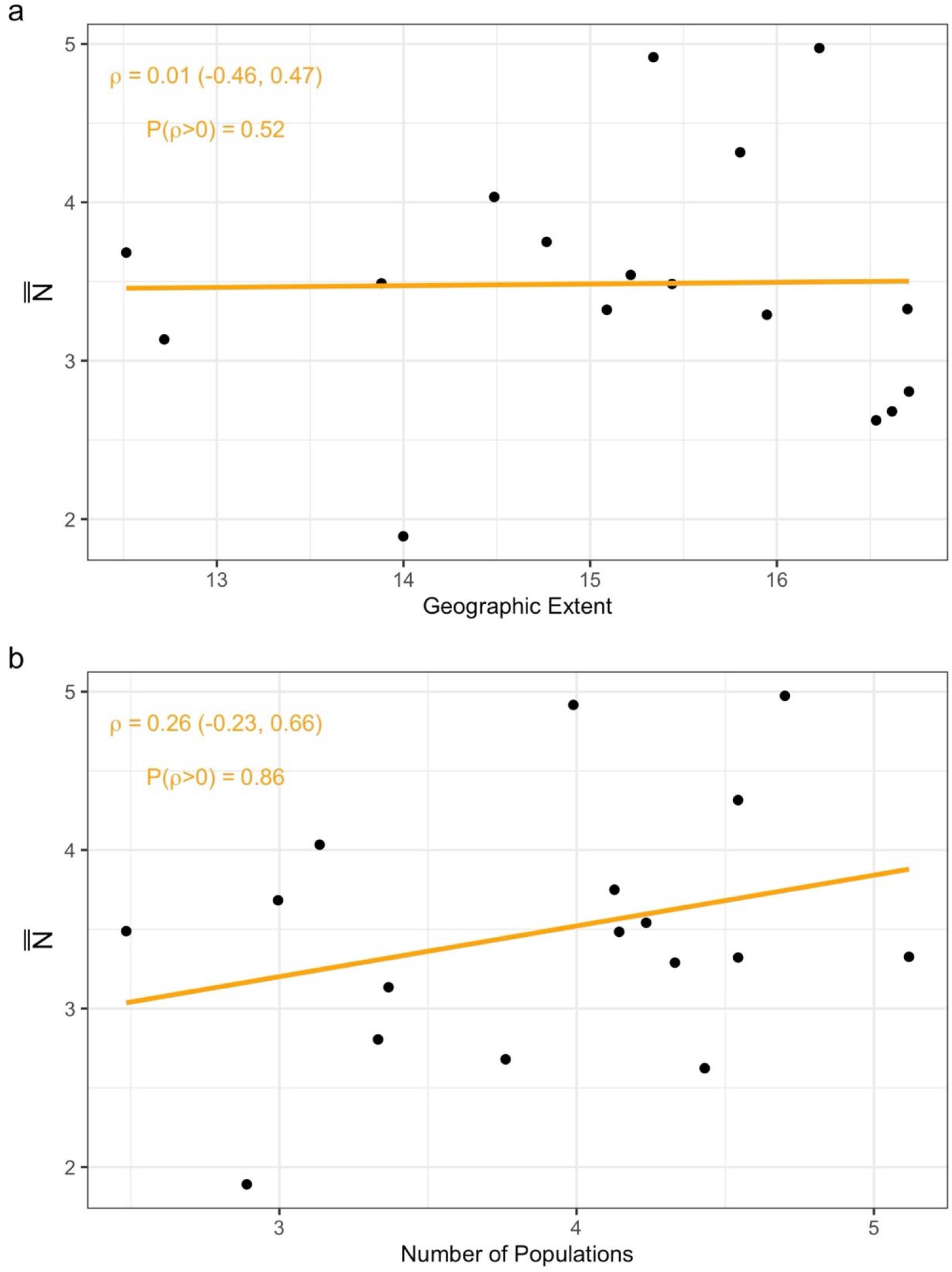
Correlations between species level median abundance 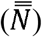 and metrics of occupancy. Every point represents a single species. The orange line is the best fit OLS line to indicate the trend in the data. *ρ* is the Pearson’s correlation coefficient and its 95% credible interval is in parenthesis. *P(ρ* > 0) indicates the probability that there is positive correlation between two parameters.

**Figure 2.**
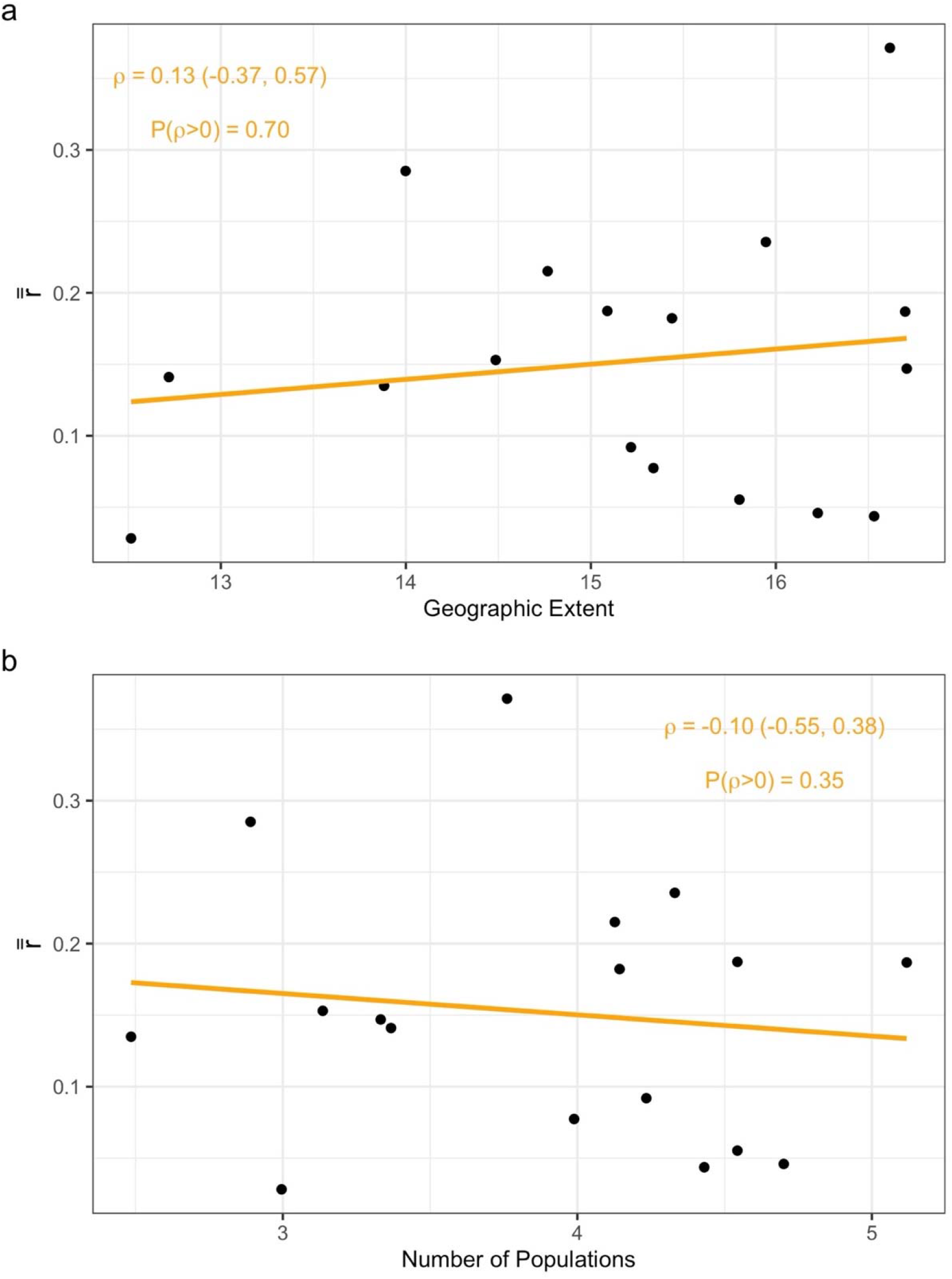
Correlations between species level median intrinsic growth rate 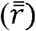 and metrics of occupancy. Every point represents a single species. The orange line is the best fit OLS line to indicate the trend in the data. *ρ* is the Pearson’s correlation coefficient and its 95% credible interval is in parenthesis. *P(ρ* > 0) indicates the probability that there is positive correlation between two parameters.

There was a positive relationship between population-level intrinsic growth rate 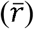 and population abundance 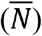 for 13 out of 17 species. The strength of these relationships were generally weak (Fig. 3). Seven species have *P(ρ* > 0) > 0.9, indicating strong evidence for the inferred positive relationship between intrinsic growth rate and abundance at the population level. Downy Woodpecker has the highest correlation coefficient with the lowest uncertainty *(ρ =* 0.42 (0.31 – 0.53)). Wrentit is the only species with a strong negative relationship between 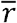 and 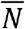 (Fig. 3).

**Figure 3.**
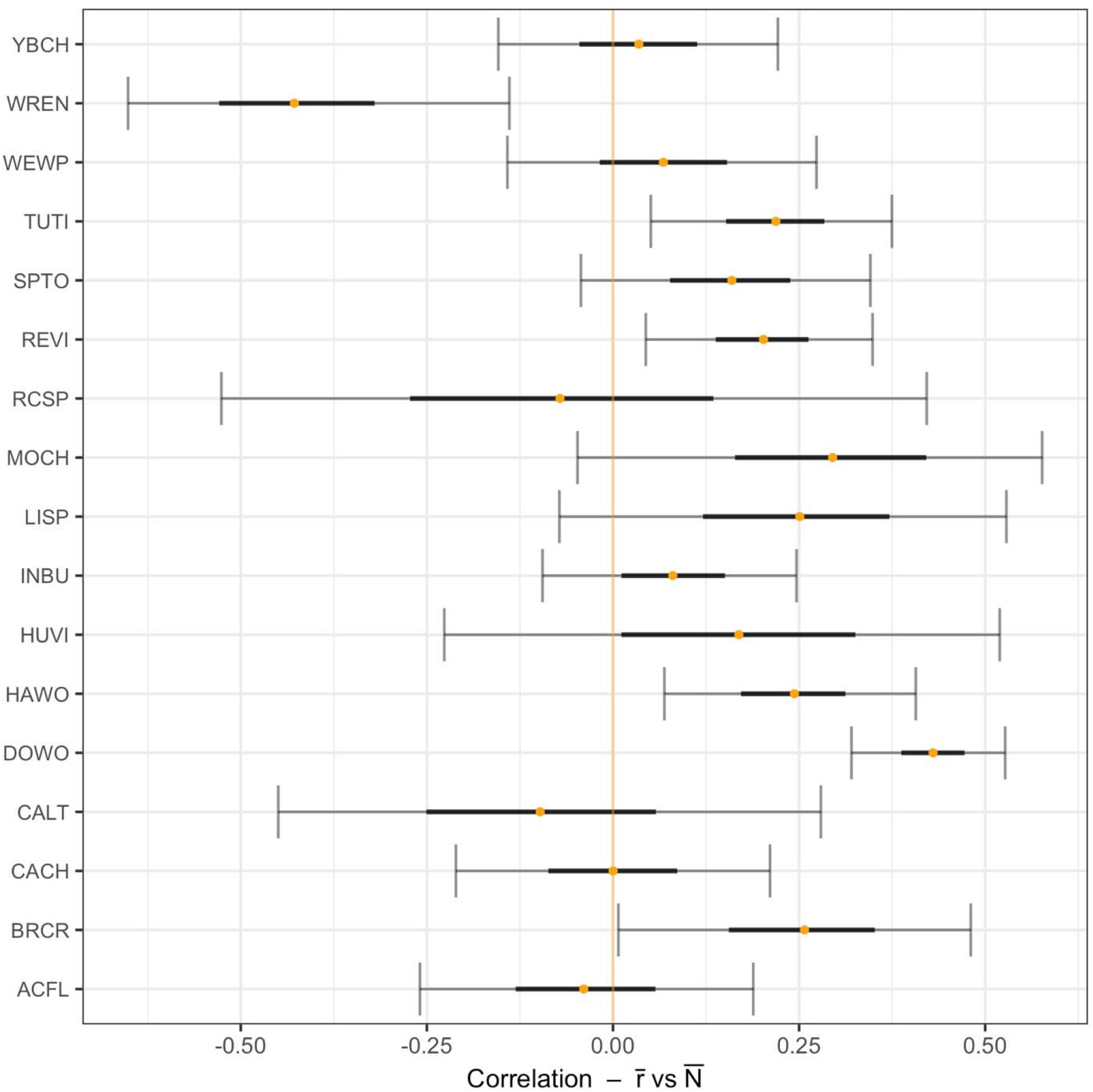
Correlation between population level abundance 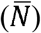 and intrinsic growth rate 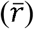 for each species. The orange dot indicates the mean of the posterior distribution of Pearson’s correlation coefficient (*ρ*). Black lines are the 50% credible intervals and gray lines are 90% credible intervals of the posterior distributions.

We found a strong negative correlation between 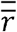 and 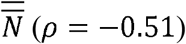 indicating a very low probability of a positive relationship (P(*ρ* > 0) = 0.01; Fig. 4). There was a strongly negative correlation between density dependence strength in survival (*β*) and 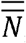, and strongly positive correlation between *β* and 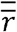, which shows that species with small average population size and large intrinsic growth rate are associated with stronger density dependence in survival (Figs. 5a and 5c). Relationship between density dependence in fecundity (*ζ*) and 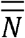 and 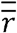 shows the opposite trend to their survival counter parts, albeit with weaker *ρ* estimates (Figs. 5b and 5d).

**Figure 4.**
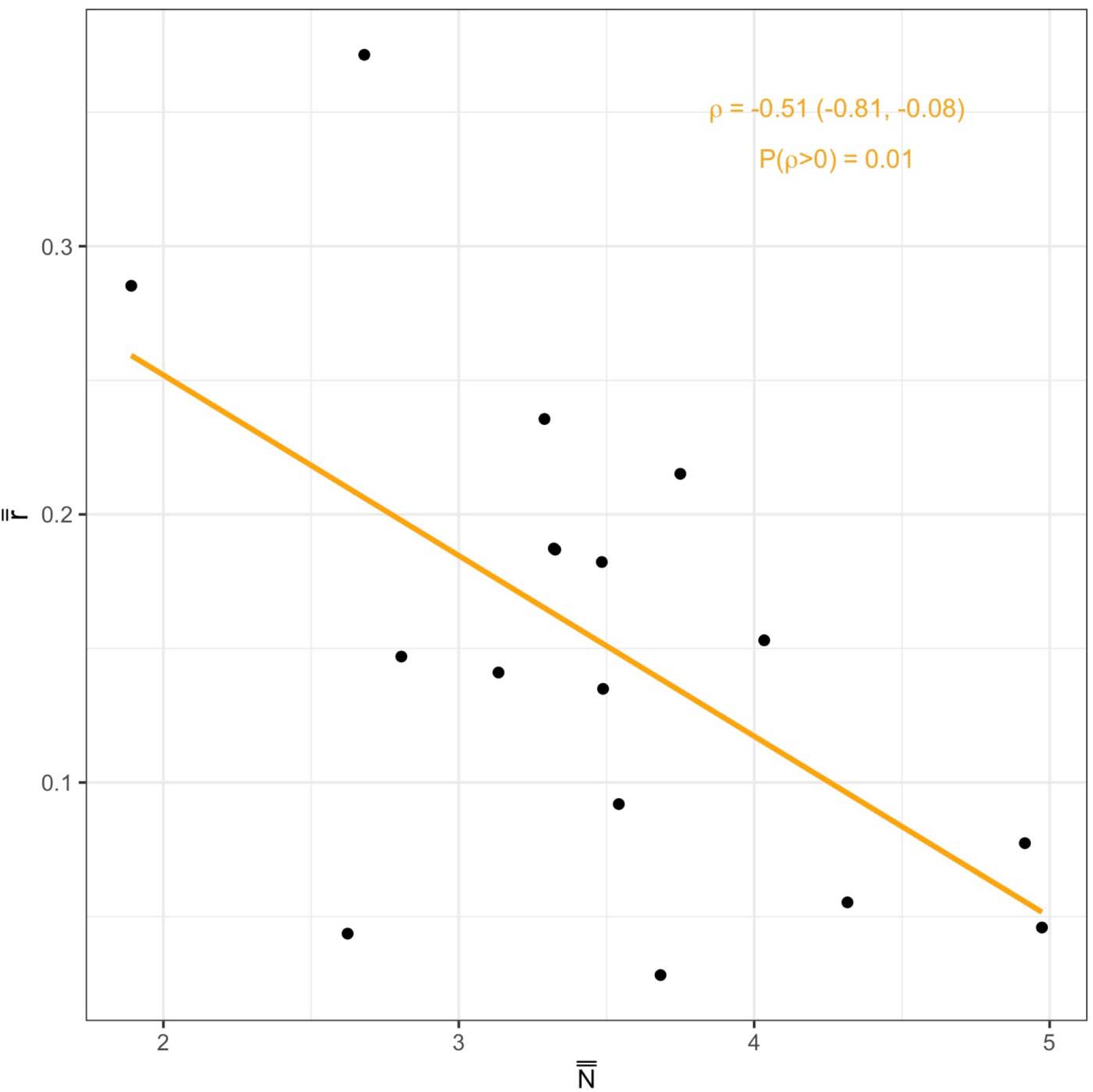
Correlation between species level median abundance 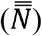, median intrinsic growth rate 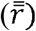. Every point represents a single species. The orange line is the best fit OLS line to indicate the trend in the data. *ρ* is Pearson’s correlation coefficient and its 95% credible interval is in parenthesis. *P(ρ* > 0) indicates the probability that there is positive correlation between two variables.

**Figure 5.**
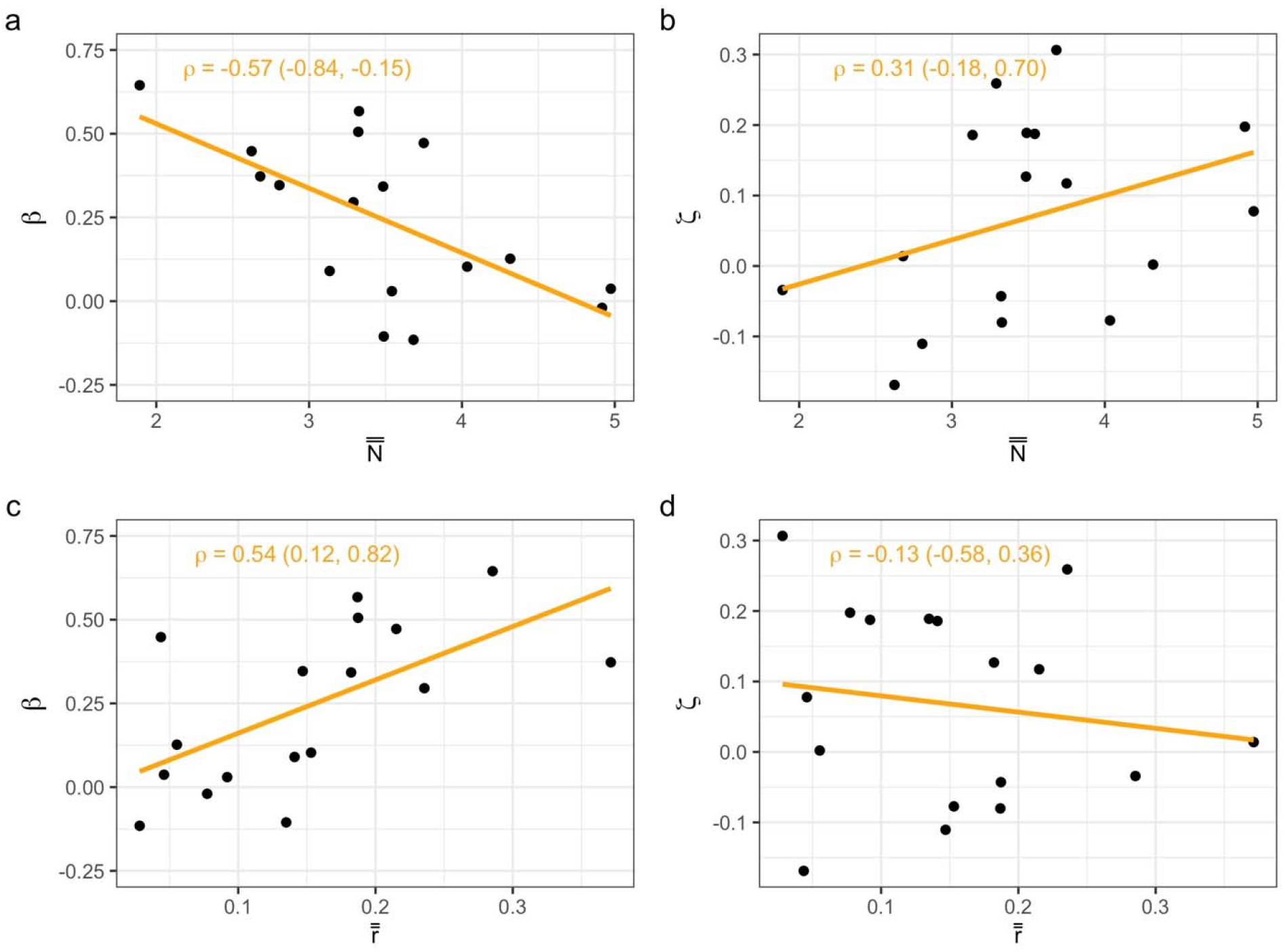
Correlations between density dependence strength in survival (*β*), density dependence strength in fecundity (*ζ*), species level median abundance 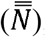, and median intrinsic growth rate 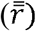. Every point represents a single species. The orange line is the best fit OLS line to indicate the trend in the data. *ρ* is Pearson’s correlation coefficient and its 95% credible interval is in parenthesis.

## Discussion

The vital rates mechanism explicitly links intrinsic population growth and density dependent population regulation with the emerging relationship between abundance and occupancy on a macroecological scale. In doing so, however, it not only provides a mechanistic, population-level explanation for the emergence of the widely reported positive abundance-occupancy relationships, but also predicts conditions under which occupancy and abundance will be unrelated. Here, we show that a group of 17 bird species have a weakly positive abundance-occupancy relationship (Fig. 1), and that this weak relationship might be caused by the fact that intrinsic growth rate is not acting as an intermediary between occupancy and abundance, as demonstrated by three lines of evidence: 1) population growth rates of these species appear mostly unrelated to their range sizes and occupancy (Fig 2), 2) intrinsic growth rates of these species appear weakly related to their abundances at the population level (Fig 3), and 3) there is a strongly negative relationship between species-level growth rate and species-level abundance (Fig. 4) which is also associated with, and potentially emerge from, differences in density dependence strengths of vital rates (Fig 5). We discuss these relationships in detail below.

A trade-off between intrinsic population growth and abundance is expected if fast growth is associated with stronger density dependence (Holt *et al*., 1997). According to the vital rates mechanism, this trade-off weakens a positive abundance-occupancy relationship because higher growth rates do not lead to higher abundances. Among the 17 bird species analyzed in this study, the trade-off between intrinsic growth rate and abundance exists among species (Fig 4). While there is no such trade-off at the population level (Fig. 3), the correlation strength between growth and abundance is not strong enough to conclude that higher growth leads to higher population abundances. There is also evidence that this species-level trade-off is driven by density-dependent population regulation because stronger density dependence is associated with higher growth and lower abundance (Fig 5). The framework we used estimates density dependence strengths for survival and fecundity separately (Ryu *et al*., 2016), which revealed subtler patterns of density-dependent population regulation compared to Holt *et al*. (1997), which used a single density dependence effect on population growth. Density dependence of fecundity and survival show opposing patterns in their relationships with abundance and intrinsic growth rate (Fig. 5). Species with lower growth rates have higher abundances and are strongly regulated by density-dependent processes, especially acting on survival rates, and species with higher growth rate, on average, have lower abundances and are more weakly regulated by density-dependent processes especially acting on fecundity (Fig. 5). We are not aware of any other study exploring these patterns, so a better understanding of the relationship of density dependence with other demographic parameters, and the macroecological patterns they might cause, requires further research of these patterns in other taxonomic groups.

While we reported weakly positive relationships between abundance and intrinsic growth rate (Fig. 3), it is important to note that this is not the relationship between annual growth and abundance but rather between their long-term averages in a given population. Holt *et al*. (1997) hypothesized that equilibrium abundance of a population (e.g. carrying capacity) is directly determined by average intrinsic growth rate but if the average abundances we estimated here (Eq. 9) do not represent equilibrium abundances, then this might explain the weak relationship with intrinsic growth rate. Time series length across populations of the 17 species varied between 5 to 17 years. It is likely that populations with the shorter time series are not in equilibrium. This creates a temporal dimension to abundance-occupancy relationships, where it is more likely to detect positive relationships for species with longer time series data if such a relationship exists (Brook & Bradshaw, 2006). The weak association we found between population level abundance and intrinsic growth rate could also indicate that they are affected by different environmental processes and can vary independently. For example, habitat loss and fragmentation can have direct impact on the size of a metapopulation without affecting the intrinsic growth rate of a species. This mismatch between processes that affect equilibrium abundance vs intrinsic growth can impact the strength of abundance-occupancy relationships, but we are not aware of any study that explicitly made this comparison.

There are several processes that can lead to a weak association between intrinsic growth rate and occupancy (Fig 2). For example, if source-sink dynamics is the dominant process determining occupancy, then it is expected that many populations with negative intrinsic growth will be occupied and occupancy will not be strongly related to population growth. We don’t have evidence to believe this is the main process in effect for these 17 bird species, however, because majority of their populations have positive intrinsic growth (Appendix S1: Fig. S1b). Another process that can weaken the relationship between occupancy and intrinsic growth is local adaptation in range-restricted species as observed in mountainous bird species in the afro-tropical region (Reif *et al*., 2006; Symonds & Johnson, 2006; Ferenc *et al*., 2016; Freeman, 2019). These mountainous species are highly adapted to high elevation areas and can reach abundance levels and population growth similar to lower elevation species while still occurring across a narrower range. Although this mechanism can explain the lack of a positive abundance-occupancy relationship of specialist species with narrow ranges, it is not applicable to our study because the majority of the17 bird species we modeled are wide-ranging across the US.

The spatial response of intrinsic growth rate to environmental factors can be different among species. Holt *et al*. (1997) hypothesized that this difference will weaken the abundance-occupancy relationship because intrinsic growth rate will lose its positive association with occupancy. They present a simple case, where one species has a relatively steep response curve and its maximum growth can be high but it is also limited to a narrow range of an environmental gradient, whereas another species has a flatter response curve that reaches a lower maximum but it can exist on a wider range of the same environmental gradient. In this example, species with the higher intrinsic growth will not have higher occupancy. In our analysis, spatial response of intrinsic growth is quantified in the slopes estimated for the environmental variables in our framework (Eq. 1a and Eq. 2a). There are two complications for comparing these slopes: 1) Because we use the principal component dimensions of environmental variables as covariates, the fitted slopes do not represent the response to same environmental variables; each dimension of each species can be different from others; and 2) The response to an environmental gradient that is described by Holt *et al*. (1997) is more akin to the concept of fundamental niche (Peterson, 2011) but the framework we used here, or any other statistical method, may not be able to estimate the “true” response to environmental variability for the simple fact that species may not occur in every suitable site because of competitive exclusion. Inter-species biotic interactions are effectively missing from the vital rates mechanism. Differences in spatial variability of intrinsic growth rates among species, and whether these differences are mainly caused by density-independent factors or biotic interactions is an important future direction for abundance-occupancy relationships research.

Even though we found a weakly positive abundance-occupancy relationship among 17 bird species (Fig 1b), we are not making any generalizations about the prevalence of positive abundance-occupancy relationships across different taxa. Detecting a positive abundance occupancy relationship depends on the selected group of species; for example, Hurlbert & White (2007) found a pre-dominantly positive abundance-occupancy relationship across 298 bird species in US but Novosolov *et al*. (2017) found no apparent pattern in 893 species across different biogeographical realms. However, uncovering mechanisms and processes associated with abundance-occupancy patterns is just as important as determining the prevalence of these patterns across multiple taxa. We believe that small sub-groups of species, such as the 17 bird species in this study, can be used to explore the processes that affect the emergence of macroecological patterns. One limitation would be the length of the time series. For example, most species in the MAPS program do not have enough recaptures to estimate data-sensitive demographic parameters such as intrinsic growth rate and density dependence. Also, long time series are necessary to detect the negative density dependence on population growth from population trends (Brook & Bradshaw, 2006). As more data become available, these demographic parameters can be estimated for more species, and these species can be divided into further sub-groups that represent different distributions of demographic parameters among them. These sub-groups would provide a more complete picture for testing the predictions of vital rates mechanism. For example, a group that has similar density dependence strength should show a stronger positive abundance-occupancy relationship compared to another group with more variable density dependence strengths. Violation of this expected pattern would indicate other factors than vital rates determining abundance-occupancy relationships.

Patterns of distribution of life on Earth are interesting in themselves, but even more so when they are associated with, and explained by, mechanisms at different levels of biological organization. Holt *et al*., (1997) presented one such mechanism for the widely reported positive abundance-occupancy relationships. It is, however, likely that the abundance-occupancy patterns are simultaneously determined by dispersal, inter-specific and intra-specific interactions, species demography and response to environmental gradients, as well as the sampling schemes used to explore these relationships. Here, we only demonstrated the effect of within-population processes. Ideally, macroecological patterns would be explored in frameworks that include dynamics at all relevant scales, including processes that are at the population, metapopulation, and assemblage or community levels, and different sampling schemes. Such cross-scale frameworks would help us get a clearer picture of the conditions under which abundance-occupancy and other macroecological relationships emerge

## Acknowledgements

We thank Kevin Shoemaker for his valuable input. This study was initiated under funding from the NASA Biodiversity Program (NNH10ZDA001N-BIOCLIM). We thank the many volunteers who have contributed to the MAPS program and the Institute for Bird Populations for development and active curation of the MAPS dataset.

