## Appendix S1 for "Inter-specific Variability in Demographic Processes Affects Abundance-Occupancy Relationships"

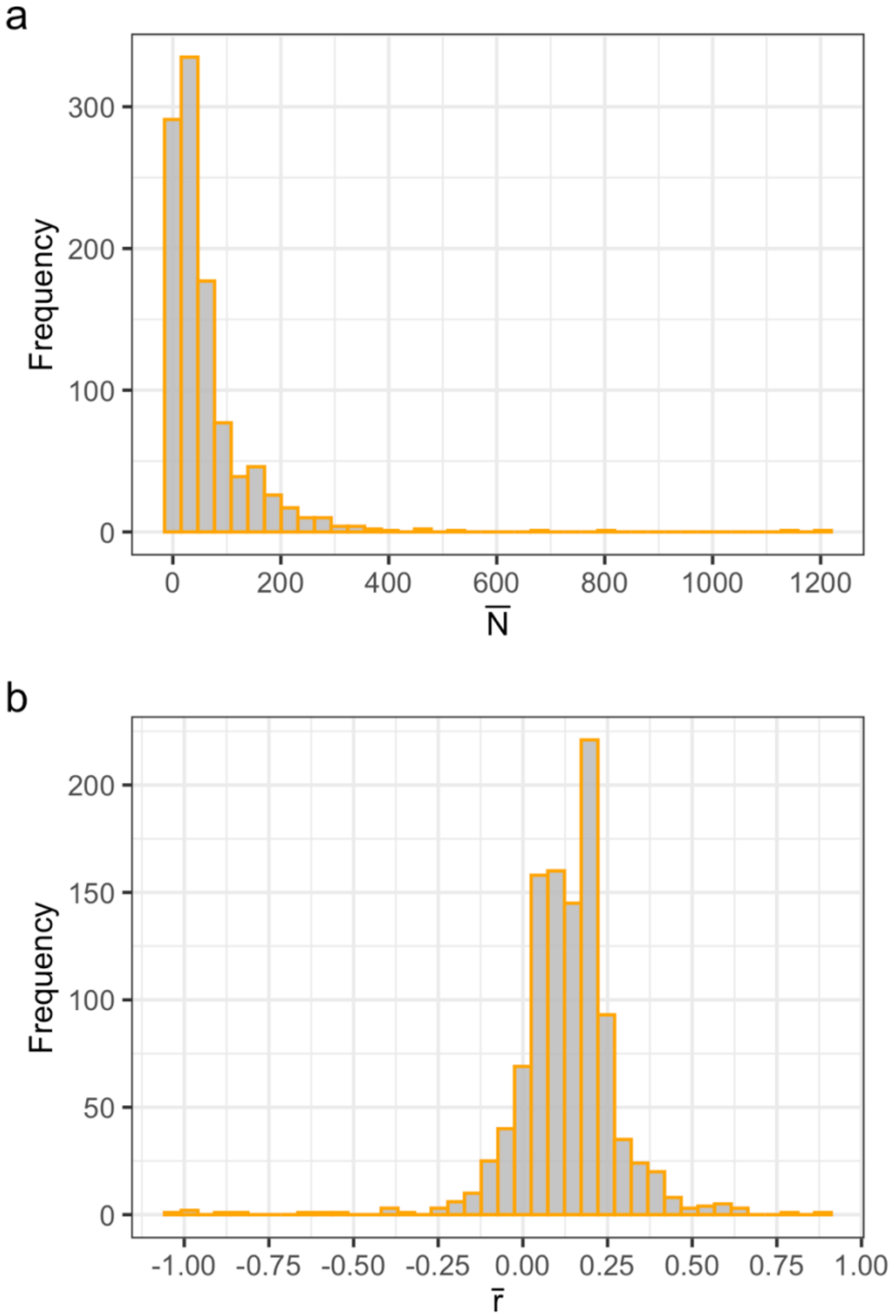


**Figure S1.** Distribution of a) population abundances ($\overline{N}$) and b) intrinsic growth rates ($\bar{r}$) across populations and species.

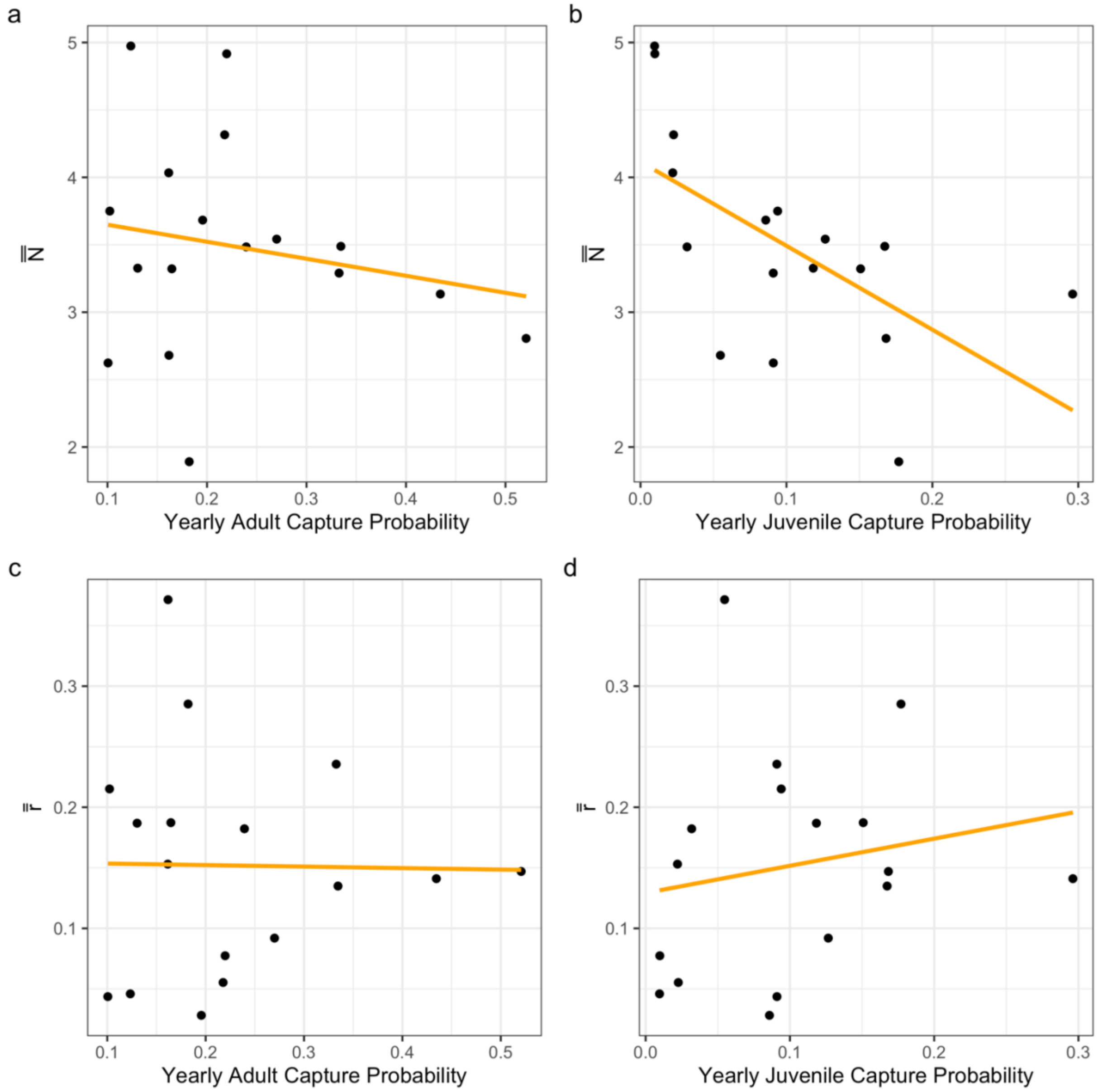


**Figure S2.** Relationship between species level median abundance ($\bar{\bar{N}}$), median intrinsic growth rate ($\bar{\bar{r}}$) and yearly capture probabilities in adult and juveniles. The orange is the best fit OLS line to indicate the trend in the data.

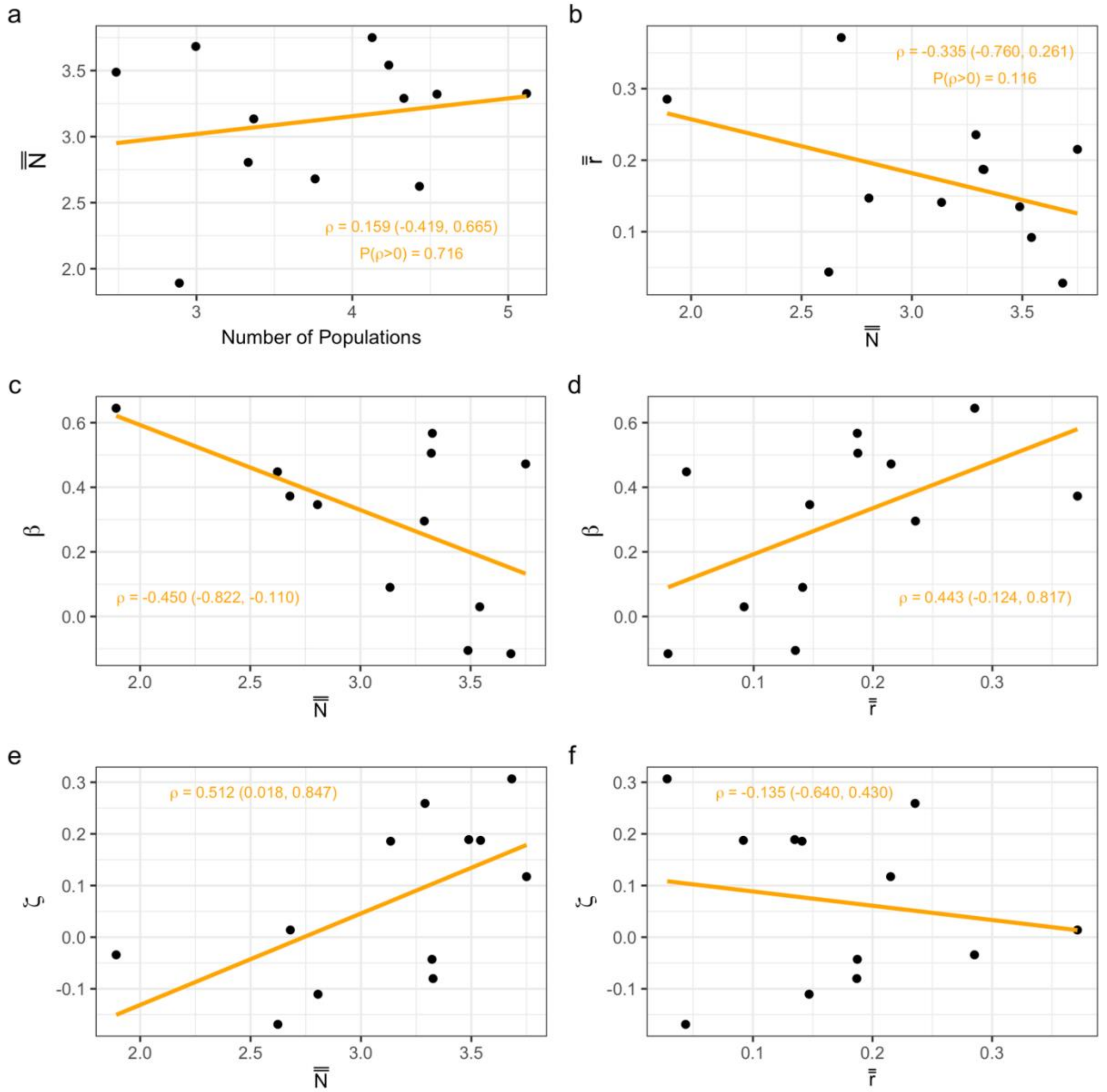


**Figure S3.** Re-drawing of some of the relationships from the figures in main text after excluding 5 species with yearly juvenile capture probabilities below 0.05.

**Table S1.** Demographic parameters as estimated by CJS-pop. 95% credible intervals are indicated in parenthesis. Survival and Fecundity estimates are reported at mean population size and at the mean values of environmental variables used in CJS-pop of a species. $\beta$ is density dependence in survival in logit scale, and $\zeta$ is density dependence in fecundity in log scale. CP is yearly capture probability and is reported at mean capture effort. $\beta$ and $\zeta$ reported here correspond to the raw output of CJS-pop. In the main manuscript and in the figures we use $-\beta$ and $-\zeta$ for easier comparison with Holt et al. (1997).

| 4-Letter Species Code | Adult Survival | Juvenile Survival | Fecundity | $\beta$ | $\zeta$ | Adult CP | Juvenile CP |
|---|---|---|---|---|---|---|---|
| ACFL | 0.574 (0.531, 0.619) | 0.309 (0.08, 0.722) | 1.902 (0.585, 5.310) | 0.020 (-0.325, 0.365) | -0.198 (-0.47, 0.068) | 0.220 (0.206, 0.233) | 0.010 (0.003, 0.027) |
| BRCR | 0.386 (0.283, 0.764) | 0.338 (0.199, 0.544) | 1.940 (0.451, 3.128) | -0.373 (-0.917, 0.343) | -0.014 (-0.177, 0.133) | 0.162 (0.037, 0.198) | 0.055 (0.036, 0.079) |
| CACH | 0.561 (0.502, 0.627) | 0.650 (0.502, 0.809) | 0.684 (0.541, 0.862) | -0.472 (-0.901, -0.010) | -0.117 (-0.24, 0.004) | 0.102 (0.090, 0.116) | 0.094 (0.078, 0.112) |
| CALT | 0.687 (0.61, 0.764) | 0.334 (0.199, 0.526) | 0.989 (0.596, 1.568) | 0.115 (-0.561, 0.849) | -0.307 (-0.619, 0.008) | 0.196 (0.171, 0.222) | 0.086 (0.054, 0.129) |
| DOWO | 0.593 (0.519, 0.688) | 0.543 (0.438, 0.656) | 0.751 (0.626, 0.892) | -0.567 (-0.843, -0.290) | 0.080 (-0.011, 0.171) | 0.130 (0.118, 0.143) | 0.118 (0.103, 0.135) |
| HAWO | 0.822 (0.719, 0.903) | 0.612 (0.391, 0.836) | 0.294 (0.176, 0.453) | -0.448 (-1.21, 0.382) | 0.169 (-0.103, 0.437) | 0.101 (0.083, 0.120) | 0.091 (0.060, 0.134) |
| HUVI | 0.601 (0.465, 0.735) | 0.368 (0.235, 0.548) | 1.113 (0.742, 1.596) | -0.645 (-1.457, 0.229) | 0.034 (-0.173, 0.223) | 0.182 (0.138, 0.232) | 0.177 (0.131, 0.233) |
| INBU | 0.533 (0.483, 0.589) | 0.669 (0.37, 0.96) | 0.744 (0.483, 1.268) | -0.127 (-0.336, 0.080) | -0.002 (-0.198, 0.196) | 0.218 (0.207, 0.228) | 0.023 (0.013, 0.035) |
| LISP | 0.505 (0.430, 0.606) | 0.362 (0.277, 0.45) | 1.384 (1.078, 1.786) | -0.346 (-0.565, -0.136) | 0.110 (0.003, 0.213) | 0.521 (0.505, 0.537) | 0.168 (0.138, 0.204) |
| MOCH | 0.504 (0.411, 0.601) | 0.126 (0.067, 0.216) | 4.286 (2.293, 7.395) | -0.103 (-0.388, 0.188) | 0.077 (-0.020, 0.174) | 0.162 (0.143, 0.182) | 0.022 (0.012, 0.038) |
| RCSP | 0.614 (0.459, 0.776) | 0.673 (0.413, 0.934) | 0.595 (0.324, 0.985) | 0.105 (-0.803, 1.233) | -0.189 (-0.508, 0.114) | 0.335 (0.282, 0.391) | 0.167 (0.104, 0.252) |
| REVI | 0.664 (0.611, 0.725) | 0.780 (0.428, 0.99) | 0.454 (0.309, 0.804) | -0.037 (-0.353, 0.300) | -0.078 (-0.414, 0.240) | 0.123 (0.115, 0.133) | 0.01 0(0.005, 0.015) |
| SPTO | 0.574 (0.528, 0.642) | 0.287 (0.223, 0.354) | 1.503 (1.187, 1.950) | -0.030 (-0.239, 0.180) | -0.187 (-0.275, -0.101) | 0.270 (0.257, 0.283) | 0.127 (0.109, 0.146) |
| TUTI | 0.500 (0.453, 0.545) | 0.519 (0.451, 0.596) | 0.967 (0.841, 1.105) | -0.506 (-0.746, -0.267) | 0.043 (-0.038, 0.125) | 0.165 (0.152, 0.178) | 0.151 (0.136, 0.167) |
| WEWP | 0.554 (0.471, 0.668) | 0.783 (0.358, 0.991) | 0.612 (0.378, 1.288) | -0.342 (-0.728, 0.047) | -0.127 (-0.444, 0.190) | 0.239 (0.221, 0.259) | 0.032 (0.014, 0.053) |
| WREN | 0.597 (0.560, 0.635) | 0.245 (0.213, 0.279) | 1.649 (1.397, 1.950) | -0.090 (-0.288, 0.106) | -0.186 (-0.264, -0.110) | 0.435 (0.419, 0.450) | 0.296 (0.275, 0.318) |
| YBCH | 0.616 (0.535, 0.731) | 0.472 (0.349, 0.599) | 0.825 (0.588, 1.140) | -0.295 (-0.455, -0.136) | -0.259 (-0.376, -0.144) | 0.333 (0.322, 0.344) | 0.091 (0.069, 0.118) |

**Table S2.** Estimates of species level median abundance ($\bar{\bar{N}}$), median intrinsic growth rate ($\bar{\bar{r}}$) and metrics of occupancy. Extent of distribution is km2 on log scale.

| 4-Letter Species Code | $\bar{\bar{r}}$ | $\bar{\bar{N}}$ | Number of populations | Number of Stations | Extent of Distribution |
|---|---|---|---|---|---|
| ACFL | 0.08 | 136.49 | 54 | 166 | 15.34 |
| BRCR | 0.37 | 14.59 | 43 | 234 | 16.62 |
| CACH | 0.22 | 42.51 | 62 | 187 | 14.77 |
| CALT | 0.03 | 39.76 | 20 | 57 | 12.51 |
| DOWO | 0.19 | 27.84 | 167 | 522 | 16.7 |
| HAWO | 0.04 | 13.79 | 84 | 364 | 16.53 |
| HUVI | 0.29 | 6.63 | 18 | 107 | 14 |
| INBU | 0.06 | 74.86 | 94 | 241 | 15.8 |
| LISP | 0.15 | 16.53 | 28 | 143 | 16.71 |
| MOCH | 0.15 | 56.48 | 23 | 141 | 14.49 |
| RCSP | 0.13 | 32.73 | 12 | 39 | 13.88 |
| REVI | 0.05 | 144.6 | 110 | 287 | 16.23 |
| SPTO | 0.09 | 34.52 | 69 | 227 | 15.22 |
| TUTI | 0.19 | 27.71 | 94 | 250 | 15.09 |
| WEWP | 0.18 | 32.6 | 63 | 223 | 15.44 |
| WREN | 0.14 | 22.97 | 29 | 121 | 12.72 |
| YBCH | 0.24 | 26.84 | 76 | 241 | 15.95 |
