## Appendix S1 for "Inter-specific Variability in Demographic Processes Affects Abundance-Occupancy Relationships"

### Appendix S2

1. CJS-pop

2. Pearson's Correlation

3. Density Dependence vs Statistical Artefacts

4. References

#### 1. CJS-pop

Here, we provide a detailed overview of CJS-pop as presented in Şen and Akçakaya (2020) with slight modifications.

CJS-pop requires, at minimum, two types of data: robust-design capture histories, and the number of captured adults and juveniles, which can be obtained from said capture histories.

1) *Capture History*: CJS-pop employs robust-design capture histories (Kendall et al. 1995) where there are multiple secondary capture occasions in a primary capture period. For example, in the test dataset we used (described later), every year between 1992 and 2008 is a primary capture period, and each month in the breeding season from May to August in each year constitute the secondary occasions. Capture of a marked individual in a secondary occasion is assigned 1, and failure of capture is assigned 0. Robust design combines open and closed population mark-recapture models. Within a primary period (here, the breeding season) populations are assumed to be closed (no mortality and no emigrations) and capture probability is estimated with captures between secondary occasions (months). This capture probability can then be used to estimate population size. Between primary periods populations are assumed to be open (individuals can die or leave the population). Survival is estimated with capture information obtained from

primary periods (for example, if an individual is captured at least once in a primary period we know it survived from the previous primary period).

2) *Number of captured adults and juveniles*: We count the number of adults and juveniles captured or recaptured at least once per year in each population. Using these counts, we estimate a population-level density index, as well as fecundity (as number of juveniles per adult). Here, we group mist-netting stations into separate populations. While we only considered a two-stage population structure in this study, the framework we present can be applied to other types of age and stage structures in populations

A survival rate estimated with CJS models is called apparent survival, because it is the joint probability of surviving and staying in the population. Apparent survival rates tend to be lower than true survival rates and may lead to biased projections if used in population models. Here, we use an additional data type to explicitly model resident and transient individuals to correct this potential bias. This is not a mandatory part of CJS-pop and can be omitted if there are not many suspected transient individuals of a modeled species. The residency model uses pre-determined residents as the third dataset of CJS-pop presented here:

3) *Pre-determined Residents*: This is a binary variable. A captured individual is categorized as a pre-determined resident if it was re-captured at least once >10 days after following its first capture in its initial year of capture. The rest of the individuals are categorized as potential transients. Transient adult individuals are assumed to be passing through the population (they will not stay and breed); transient juvenile individuals are assumed to leave the population after fledging. Residents categorized this way are named pre-determined, because their categorization happens before running any model (Saracco et

al. 2010). Pre-determined residents are assigned 1 and potential transients 0 in this data set.

These three data sets, derived from the same robust-design capture histories, are used to build 5 connected models and estimate their associated parameters: Survival, Capture, Density, Residency, and Fecundity.

*Survival*

Survival is modeled as a function of breeding stage (juvenile or adult), and a population density index:

$$\text{logit}(\phi_{x,k,t}) = \alpha_x - \beta_1 \cdot D_{k,t} + \beta_2 \cdot W_{1,k,t} + \beta_3 \cdot W_{2,k,t} + \epsilon_t \qquad (1a)$$

$$\epsilon_t \sim \text{Normal}(0, \sigma_s^2) \qquad (1b)$$

Where, $x = 1,2, \dots, X$; $k = 1,2,3\dots, K$; $t = 1,2,3\dots, T$; $\phi_{x,k,t}$ is the survival probability of a stage $x$ invidiual, at population $k$, and at year $t$; $\alpha_x$ is the survival probability of a stage $x$ individual in logit scale at mean population size and at mean of environmental covariates $W_1$ and $W_2$; $\beta_1$ is the change in survival in logit scale with one unit change in population density index; $D$ is the population density index at population $k$, and at year $t$; $\beta_2$ and $\beta_3$ are the change in survival in logit scale with one unit change in $W_1$ and $W_2$, respectively; $\epsilon_t$ is the temporal random effect at year $t$, and $\sigma_s^2$ is the temporal process variance of survival at logit scale, which assumes full correlation between populations in their temporal variability (all populations have the same temporal effect each year).

*Capture*

Capture probability is modeled as a function of effort and breeding stage:

$$\text{logit}(p_{x,k,t,h}) = \gamma_x + \delta \cdot E_{k,t,h} \qquad (2)$$

where, $h = 1,2,3\dots, H$; $p_{x,k,t,h}$ is capture probability of a stage $x$ individual at population $k$, year $t$, and month $h$; $\gamma_x$ is the monthly capture probability on logit scale of a stage $x$ individual at mean capture effort; $\delta$ is the change in monthly capture probability on logit scale with one unit change in capture effort; and $E$ is the capture effort.

*Density*

The monthly capture probabilities obtained from the capture model are used to calculate the yearly capture probabilities, and these in turn are used to estimate the expected size of each population in each year. Yearly capture probability for each adult and juvenile in a given year and population is calculated as follows:

$$P_{x,k,t} = 1 - \prod_{h=1}^{4}\left(1 - p_{x,k,t,h}\right) \qquad (3a)$$

where, $P_{x,k,t}$ is the probability that a stage $x$ individual will be captured at least once in 4 months of the breeding period at year $t$ and population $k$.

Using the heuristic estimator for population size (first term on the right-hand side of equation 3b) with a correction for years with 0 captures (Dail and Madsen 2011; second term on the right-hand side of equation 3b), the numbers of adults and juveniles can be derived as:

$$N_{x,k,t} = \frac{n_{x,k,t}}{P_{x,k,t}} + \frac{\left(1 - P_{x,k,t}\right)}{P_{x,k,t}} \qquad (3b)$$

where, $n_{x,k,t}$ is the number of captured stage $x$ individuals in population $k$ and year $t$, and $N_{x,k,t}$ is the expected number of stage $x$ individuals of the same population and year combination.

Expected total population size of population $k$ at year $t$, $M_{k,t}$, is estimated by adding up $N_{1,k,t}$ and $N_{2,k,t}$. Density index of population $k$ at year $t$, then, is estimated as:

$$D_{k,t} = \frac{M_{k,t}}{\frac{\sum_{t=1}^{T} M_{k,t}}{T}} - 1 \tag{4}$$

where, $T$ is the number of years with capture effort in population $k$. Because relative density is calculated by dividing each year's population size (numerator in equation 4) by the average size of that population across years (denominator in equation 4), average relative density of a given population is always 1. When modelling survival and fecundity, we use this information to centralize the relative density estimate around its mean by subtracting 1 and refer to this metric as population density index ($D$). Centralized nature of this metric increases the effective number of samples in the density dependence strength parameters estimated by survival and fecundity models ($\beta$ and $\zeta$, respectively).

*State and Observation Processes*

We model survival and capture probabilities as functions of population-level covariates (population density index and effort, respectively). However, capture history data are at the individual level, so there needs to be a link between population-level parameters and individual-level data. We make this link by using the matrix $S_{i,t}$ to indicate the breeding stage of the *i*th individual in year $t$, and the vector $g_i$ to indicate the population that the *i*th individual is in. The state of the *i*th individual (alive and in the population or not) in year $t + 1$, $Z_{i,(t+1)}$, depends on whether it was alive at time $t$, whether it survived the period from $t$ to $t + 1$, and whether it is a resident. This latent state is a Bernoulli random variable:

$$Z_{i,(t+1)} \sim \text{Bernoulli}\left(Z_{i,t} \cdot \phi_{(S_{i,t}),(g_i),t} \cdot R_i\right) \tag{5}$$

where, $\phi_{(S_{i,t}),(g_i),t}$ is the survival probability at the breeding stage and population of the $i$th individual from year $t$ to $t + 1$, and $R_i$ is the true residency status of the *i*th individual (1 if it is a resident, 0 if it is a transient).

Every element of the capture history, $y_{i,t,h}$ (1 if an individual is captured, 0 if it is not), is a Bernoulli random variable with a monthly capture probability, $p_{(S_{i,t}),(g_i),t,h}$, conditional on the individual being alive and in the population, $Z_{i,t}$.

$$y_{i,t,h} \sim \text{Bernoulli}\left(Z_{i,t} \cdot p_{(S_{i,t}),(a_i),t,h}\right) \tag{6}$$

*Residency*

We employ a simplified version of the models described in Saracco et al. (2010) when modelling residents. This approach accounts for transients that will leave the population after their first capture but does not model movement between populations. We assume that there are two parameters, $\rho_1$ for juveniles and $\rho_2$ for adults, that governs the probability that we can correctly categorize a true resident as a pre-determined resident. The residency data (1 for pre-determined residents, 0 for potential transients), $r_i$, is a Bernoulli random variable and is conditional on the $i$th individual being a true resident ($R_i$):

$$r_i \sim \text{Bernoulli}\left(R_i \cdot \rho_{S_{i,f_i}}\right) \tag{7}$$

where, $f_i$ is the first capture year of the $i$th individual, and $S_{i,f_i}$ corresponds to the stage of the $i$th individual in its first capture year. We also assume that there is a parameter for each stage, $\pi_1$ and $\pi_2$, that governs the probability that a juvenile or an adult in the population is a true resident, respectively. $R_i$ is a Bernoulli random variable with probability $\pi_{S_{i,f_i}}$:

$$R_i \sim \text{Bernoulli}\left(\pi_{S_{i,f_i}}\right) \tag{8}$$

This model effectively filters out some of the capture histories that only has the first capture followed by no captures for the remainder of study period (a single 1 and trailing 0s in their capture history). If true residency of an individual ($R$) is estimated to be 0 (a transient), then its state ($Z$), will also be automatically 0 (see eq. 5) and it will not lower the survival estimate. $\pi$ and $\rho$ are nuisance parameters and are not used in projections explained below.

*Fecundity*

We followed Ryu et al. (2016)'s fecundity definition: the annual ratio of juveniles to adults for each defined population. Fecundity was modeled with a log link:

$$\log(F_{k,t}) = \theta - \zeta_1 \cdot D_{k,(t-1)} + \zeta_2 \cdot W_{1,k,t} + \zeta_3 \cdot W_{2,k,t} + \omega_t \qquad (9a)$$

$$\omega_t \sim Normal(0, \sigma_f^2) \qquad (9b)$$

where, $F_{k,t}$ is fecundity at population $k$ and year $t$; $\theta$ is the average fecundity in log scale at mean population size and at mean of environmental covariates $W_1$ and $W_2$; $\zeta_1$ is the change in fecundity in log scale with one unit change in population density index; $\zeta_2$ and $\zeta_3$ are the changes in fecundity in log scale with one unit change in $W_1$ and $W_2$, respectively; $\omega_t$ is the temporal random effect at year $t$, and $\sigma_f^2$ is the temporal process variance of fecundity in log scale.

This function is then linked to the observed number of juveniles as a zero inflated Poisson regression:

$$N'_{1,k,t} = N_{2,k,t} \cdot F_{k,t} \qquad (10)$$

$$n_{1,k,t} \sim Poisson(P_{1,k,t} \cdot N'_{1,k,t} \cdot w_{k,t}) \qquad (11a)$$

$$w_{k,t} \sim Bernoulli(\psi) \qquad (11b)$$

where, $N'_{1,k,t}$ is the expected number of juveniles at population $k$ and year $t$, which is estimated by multiplying the fecundity at population $k$ and year $t$ ($F_{k,t}$) with $N_{2,k,t}$ that is the expected number of adults estimated by equation 3b at population $k$ and year $t$; $\psi$ is the probability that the number of captured juveniles in a given year and population is not 0, and $w_{k,t}$ is the indicator as a Bernoulli random variable stating whether number of captured juveniles in population $k$ and year $t$ is higher than 0. In this fecundity model, we only included years and populations that had at least 1 adult capture. We used a zero-inflation model because its fecundity estimates were biologically more realistic and had Bayesian p-values closer to 0.5 compared to a regular Poisson model.

The expected number of juveniles is estimated twice in the CJS-pop framework: once as a derived variable ($N_{1,k,t}$) using the heuristic population size estimator, and a second time ($N'_{1,k,t}$) as a function of density and expected number of adults. If we use only $N'_{1,k,t}$, commonly used MCMC software (e.g. JAGS) do not run due to the cyclical nature of CJS-pop. In the current state of CJS-pop, $N_{1,k,t}$ is employed in density index estimation, and $N'_{1,k,t}$ in fecundity estimation.

*Priors*

We used weakly informative priors for all parameters in the framework. Gelman (2006) defines a prior distribution as weakly informative if it is proper and the information it provides is intentionally weaker than actual available prior knowledge. In the same vein, in terms of the information it encodes, a weakly informative prior is considered to be between objective and expert priors (Simpson et al. 2017), or structural and regularizing priors (Gelman et al. 2017). Additionally, we used the following growth rate ($\lambda$) equation to set the priors of vital rates:

$$S_A + S_J * F = 1 \tag{12}$$

where, $S_A$ and $S_J$ are average survival rates of adults and juveniles at carrying capacity and $F$ is the fecundity at carrying capacity. This equation states that population growth rate ($\lambda$) is equal to

1 at carrying capacity. The degrees of freedom for these three parameters is 2. If we know the estimates of two of these parameters, the third one can be directly calculated. Among these three, juvenile survival is the hardest to estimate because juvenile recaptures are rare. So, we only used priors for adult survival and fecundity and calculated juvenile survival as:

$$S_J = \frac{1 - S_A}{F} \tag{13}$$

In the density independent models in which we modeled only the average vital rates we assumed all three rates were independent and did not use this relationship.

The list of all priors used for CJS-pop fit to Brown Creeper data is as follows:

$$\begin{aligned}
S_{ad} &\sim U(0,1) \\
\beta &\sim t(0,10,1) \\
\sigma_s &\sim t(0,10,1)[0,\infty] \\
\gamma_{ad} &\sim t(0,10,1) \\
\gamma_{juv} &\sim t(0,10,1) \\
\delta &\sim t(0,2.5,1) \\
F &\sim N(0,10)[0,\infty] \\
\zeta &\sim t(0,5,1) \\
\sigma_f &\sim t(0,5,1)[0,\infty] \\
\pi_{ad} &\sim U(0,1) \\
\pi_{juv} &\sim U(0,1) \\
\rho_{ad} &\sim U(0,1) \\
\rho_{juv} &\sim U(0,1)
\end{aligned}$$

*t* states a t distribution where the first argument is the mean, the second is the standard deviation, and third is degrees of freedom. *N* is normal distribution where the first argument is the mean, and the second is standard deviation. *U* is the uniform distribution where arguments indicate the low and high limits of the distribution. “[,]” is used to indicate between which limits a distribution was truncated.

*Model Specifications*

We used JAGS 4.3 (Plummer, 2003) as the MCMC sampler with R packages R2Jags (Su and Yajima, 2015) and rjags (Plummer, 2019). Post-hoc data wrangling was done using the R package MCMCvis (Youngflesh 2018). We ran the models with 4 chains, 100000 iterations, 50000 burn in and a thinning rate of 20. We checked convergence with R-hat values, and assumed chains were converged when Rhat was <1.05. Rhat and effective sample size of MCMC chains is calculated by MCMCvis package, which follows Brooks and Gelman (1998). The R code for data manipulation and JAGS code for the framework is available at https://github.com/bilgecansen/CJS-pop.

**2. Pearson's Correlation**

$$\boldsymbol{Y} \sim MN(\boldsymbol{\mu}, \boldsymbol{\Sigma})$$

$$\boldsymbol{\Sigma} = \begin{bmatrix} \sigma_1^2 & \sigma_1 \cdot \sigma_2 \cdot \rho \\ \sigma_1 \cdot \sigma_2 \cdot \rho & \sigma_2^2 \end{bmatrix}$$

where, $\boldsymbol{Y}$ is a matrix with two columns for two variables and $I$ rows; $\boldsymbol{\mu}$ is the vector for populations means ($\mu_1$ and $\mu_2$) of two variables; $\boldsymbol{\Sigma}$ is the covariance matrix for two variables; $\sigma_1^2$ and $\sigma_2^2$ are variance of the first and second variables, respectively; $\rho$ is the correlation coefficient between two variables; $MN$ indicates a multivariate normal distribution.

*Priors*

$$\begin{aligned} \mu_1 &\sim N(0, 100) \\ \mu_2 &\sim N(0, 100) \\ \sigma_1 &\sim U(0, 100) \\ \sigma_2 &\sim U(0, 100) \\ \rho &\sim U(-1,1) \end{aligned}$$

*Model Specifications*

We used JAGS 4.3 (Plummer, 2003) to estimate Pearson's correlation coefficient with an MCMC sampler. We ran the sampler for 35000 iterations with three chains and thinning rate of 3 while omitting first 10000 iterations as burn in. The R code for data manipulation and JAGS code for the framework is available at https://github.com/bilgecansen/VitalRatesTest.

**3. Density Dependence and Statistical Artefacts**

Species with lower average yearly capture probability (cp) tend to be associated with higher $\bar{\bar{N}}$. This trend is more pronounced with juvenile cp (Figure S2b). To explore whether this trend introduced biases to the results reported in the main text, we removed species with a juvenile cp < 0.05 (White-Eyed Vireo, Red-Eyed Vireo, Mountain Chickadee, Arcadia Flycatcher, Indigo Bunting). After removing these species, the strongest relationships seen in Figures 1 and 4 were slightly weakened while their trends remained unchanged (Figure S3).

The negative correlation between $\bar{\bar{r}}$ and $\bar{\bar{N}}$ is driven by their relationship with density dependence strength (Figure 4). Strength of density dependence in survival ($\beta$) is negatively and positively correlated with $\bar{\bar{N}}$ and $\bar{\bar{r}}$, respectively. While strength of density dependence in fecundity ($\zeta$) is also correlated with $\bar{\bar{N}}$ and $\bar{\bar{r}}$, the correlation strength here is much weaker compared to density dependence in survival. There is no statistical mechanism stemming from CJS-pop's model structure that cause the correlation between $\beta$ and $\bar{\bar{N}}$. There is, however, a statistical component to the correlation between $\beta$ and $\bar{\bar{r}}$. CJS-pop models adult survival, juvenile survival, and fecundity in a structure in which that they co-vary (via a prior structure explained above), but it does not force any one of them to be equal among species. For example, survival and fecundity estimates could have also varied among species to the point that $\beta$ and $\zeta$ estimates would be closer, essentially meeting one of Holt *et al.*, (1997)'s fundamental assumptions. We see this partially happen in Figure 4d where the relationship between $\bar{\bar{r}}$ and $\zeta$ is weak. Because the pattern we report on Figure 4c arose independently, and not as a statistical artefact, we argue that the relationship between $\bar{\bar{r}}$ and $\beta$ also carries an ecological signal.

It is also important to note that the relationship of $\beta$ and $\zeta$ with $\overline{\overline{N}}$ and $\bar{\bar{r}}$ show opposite signs (Fig 4a vs. 4b; and 4c vs. 4d). This pattern carries a mix of statistical and ecological signals. CJS-pop uses vague priors for adult survival and fecundity and calculates juvenile survival with the assumption that at mean population density $r$ will be 0. This prior structure links vital rates and their density dependence estimates so that they don't vary independently outside the expectations of density dependent dynamics. So, a high $\beta$ might be balanced with a low $\zeta$, or low adult survival and fecundity might lead to the estimation of higher juvenile survival. However, CJS-pop does not force $\beta$ and $\zeta$ to have a negative or positive relationship with $\bar{\bar{r}}$ and $\overline{\overline{N}}$. The fact that we observe these particular trends (Figure 4) could indicate an ecological signal signifying the density dependence dynamics of the modeled species.
